# Alcohol consumption, obesity and hypertension: Relationship patterns along different age groups in Uganda

**DOI:** 10.1101/654251

**Authors:** Nazarius Mbona Tumwesigye, Gerald Mutungi, Silver Bahendeka, Ronald Wesonga, Monica H. Swahn, Agaba Katureebe, David Guwatudde

**Author notes:** Corresponding author (NMT). NMT initiated the paper and produced the first draft. All co-authours have made a great contribution in different ways.

## Abstract

**Introduction:** Uganda is experiencing a significant increase in the prevalence of non-communicable diseases including hypertension and obesity. Frequent alcohol use is also highly prevalent in Uganda and is a key risk factor for both hypertension and obesity. This study determines the trends of frequent alcohol consumption, hypertension and obesity across different age groups, and the extent to which alcohol consumption affects the two.

**Methods:** The data were extracted from the 2014 National Non-communicable Diseases Risk Factor Survey (N=3,987) conducted among adults aged 18 to 69 years. Hypertension was defined as systolic blood pressure ≥140mmHG or diastolic blood pressure ≥90. Obesity was defined as body mass index >30 kg/m^2^. Frequent alcohol consumption was defined as alcohol use 3 or more times a week. Multivariable log binomial regression analysis was carried out for each of the two outcome variables against age group and controlled for frequency of alcohol consumption and few other independent factors. Non-parametric tests were used to compare trends of prevalence ratios across age groups. Modified Poisson regression was use in few instances when the model failed to converge.

**Results:** The results showed increasing trend in the prevalence of hypertension and frequent alcohol consumption but a declining trend for obesity along different age groups (p<0.01). Frequency of alcohol consumption did not significantly modify the age group-hypertension and age group-obesity relationships although the effect was significant with ungrouped age. There was significance in difference of fitted lines for hypertension prevalence ratios between frequent drinkers and mild drinkers and between abstainers and frequent drinkers. Alcohol consumption did not have any significant effect on obesity-age group relationship.

**Conclusion:** The results call for more research to understand the effect of alcohol on the hypertension-age relationship, and the obesity-age relationship. Why prevalence ratios for hypertension decline among those who take alcohol most frequently is another issue that needs further research.

## Introduction

About 13% of the world’s adult population (11% of men and 15% of women) is obese[1] while 32% of adults aged more than 24 years are hypertensive[2]. The prevalence of obesity and obesity-related diseases including hypertension are increasing worldwide [3]. These conditions lead to reduced quality of life given their protracted nature, and they also lead to premature deaths, especially due to cardiovascular diseases and diabetes [4]. Once associated only with high income countries, obesity and hypertension are now highly prevalent in low and middle income countries, including Uganda, and are shown to be on the rise[5, 6]. The World Health Organization (WHO) projects the number of hypertension cases in Sub Sahara Africa (SSA) to increase substantially from an estimated 80 million in 2000 to 150 million in 2025[7].

Alcohol consumption is widely known to be associated with high blood pressure and obesity [5, 8–11] both of which are known to increase with age [12, 13]. However, little is known on how alcohol affects high blood pressure and obesity trajectories across age. So far studies have reported varying patterns of relationship between alcohol and hypertension across different age groups. A study in the United States of America, found that in young people aged 18-26, blood pressure reduced among those who took 2-3 drinks a day but rose higher with more or less alcohol intake [14]. In Germany a study found a linear relationship between alcohol intake and blood pressure for only men aged 20-34 and 50-74 and women aged above 49 years[15]. In France a positive relationship between blood pressure and alcohol intake was more evident in under 40 years[16]. A study in Japan found that the elevating effect of alcohol drinking on blood pressure was more prominent in the elderly than in the young[17]. A study in Netherlands found a stronger association between alcohol and blood pressure in older men compared with young men[18]. A study in Michigan, USA found that alcohol intake patterns significantly changed the relationship between age and blood pressure[19]. Such varying evidence calls for more localized research that can inform local intervention. Africa has the lowest research output on most health fields but more critically missing is evidence on Non-communicable diseases and their risk factors in the local environment [20].

In 2016, 7% of women and 1% of men in age group 15-49 in Uganda were obese [21]. In 2014, the National Non-Communicable Diseases (NCD) Risk Factor Survey found that the prevalence of hypertension was 26.4% and it was associated with Body Mass Index (BMI) > 25kg/m^2^ [6, 22]. These estimates are consistent with findings from other studies conducted in smaller populations in Uganda which estimated the prevalence of hypertension to be in the range of 14%-35% [11]. On alcohol consumption the country is rated 5^th^ highest consumer in Africa in terms of per capita pure alcohol with an estimated average consumption of 9.8 liters (14.4 liters for males and 5.2 liters for females) of pure alcohol per person per year[23].

There is a paucity of research on how alcohol affects obesity and hypertension levels across age groups [24, 25] especially in developing countries yet this would inform age group specific intervention. As such, the purpose of this study is to establish the trends of frequent alcohol consumption, hypertension and obesity across different age groups and the extent to which the frequency of alcohol consumption influences the trends of the two.

## Materials and Methods

This paper uses secondary data from the National Non-Communicable Disease (NCD) Risk Factor Survey of 2014. The survey included 3,987 participants aged 18-69 and 60% of them were females. The data were collected using the STEPwise approach to surveillance (STEPS). STEPS is a World Health Organization method that provides a standardized method for collection, analysis and dissemination on risk factors for non-communicable diseases (NCD)[22]. The survey covered the whole country and used a three stage sampling design to select participants. The first stage involved sampling enumeration areas (EA), followed by random selection of 14 households in each EA and lastly a random selection of one member of household from a list of eligible members. The response rate from the survey was 99% and more details on the methods used can be obtained from the national NCD report or papers written from the mother data set [6, 22].

For purposes of this study, only relevant variables were provided by the managers of the NCD survey. Key among these variables were frequency of alcohol consumption and amount drank in previous 30 days, height, and weight, biometrics that include hypertension and body mass index, socio-demographic-economic characteristics of the respondents.

Hypertension is defined as systolic blood pressure ≥140 mmHg or diastolic blood pressure ≥90 mmHg [26] while obesity is defined as body mass index greater than 30 kg/m^2^ [27]. Measurements for Systolic and diastolic blood pressure were taken three times and average for each was computed. In the previous study the average was computed for only last 2 measurements and the difference is minor. Frequent alcohol consumption was measured as taking alcohol 3 or more times a week.

We used log binomial regression to model hypertension and obesity with key independent variable being age group and key interaction variable as frequency of alcohol consumption. Binomial models were preferred because they provide prevalence ratios directly. Secondly unlike the alternative logistic regression log binomial models do not overestimate their coefficients when the outcome of interest is a common occurrence[28, 29] although they also have problems of lack of convergence[19]. In the few times non-convergence occurred we used modified Poisson regression which solves the problem but it is not also perfect since it produces inconsistent variances[19]. Stata V14 software was used for analysis.

Log binomial models are expressed as follows[29]

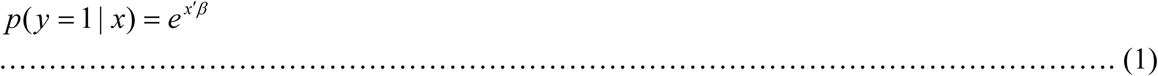

Where

*p* = Probability of occurrence of an event of interest in this study this is being hypertensive or obese
*y* = Outcome of interest. This can be 1 (occurred) or 0 (did not occur)
*x* = Covariate
*x*′*β* = *β*_0_ + *β*_1_ *x*_1_ + *β*_2_ *x*_2_ + …… + *β_n_ x_n_*. This is the model’s linear predictor where the covariates can be continuous or dichotomous.

From the above the prevalence ratio is computed as an exponentiation of the product of the coefficients and the difference in covariate values:

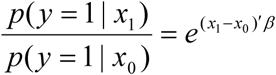

For a model with one dichotomous covariate the prevalence ratio is

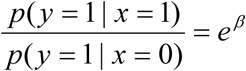

For modified Poisson regression

The following model is fitted but with robust variance estimators that will narrow the confidence intervals of the estimates[30].

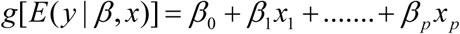

Where 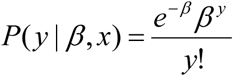

*y* = 1,2,3…

*g* = Link function

Age was the main independent variable while alcohol consumption was the main modifying factor under investigation. Charts were used to show trends of prevalence ratio for hypertension and obesity across age groups and effect of alcohol consumption.

## Ethics

The conduct of the National NCD Risk Factor Survey was approved by the Institutional Review Committee of St. Francis Hospital Nsambya, Kampala, Uganda, and registered by the Uganda National Council for Science and Technology. Written informed consent was obtained from eligible subjects before enrollment in the study. Participants with an average systolic blood pressure readings of at least 120 mm Hg, and/or diastolic blood pressure of at least 80mm Hg, reporting not to be on treatment for hypertension, were advised to as soon as possible report to the nearest government owned health facility for further evaluation. The Uganda ministry of health granted permission to use the data for this work.

## Results

### Characteristics of respondents

A total of 2956(74.1%) of the respondents were in the age range 21-50 years. The age distribution did not differ by sex (Table 1). Two thirds were married or in relationship, but marital status varied by sex. Two fifths had attained primary school, but among women a higher proportion did not have any formal education. Nearly two thirds were employed, but among men a higher percentage were employed than among women (75% vs. 58%). The median income per month was 100,000(≈USD 30) and it was significantly higher among men (110,000) (USD ≈33) than women (60,000) (≈18).

**Table 1:** Background Characteristics of the Respondents in NCD Survey

| Characteristics | Male's<br>N (%) | Female's<br>N (%) | All<br>N (%) | Chi-sq. test<br>p-value |
| --- | --- | --- | --- | --- |
| **Age** |  |  |  |  |
| 18-20 | 171(10.7) | 257(10.8) | 428(10.7) |  |
| 21-25 | 278 (17.3) | 392 (16.5) | 670 (16.8) |  |
| 26-30 | 261 (16.3) | 412 (17.3) | 673 (16.9) |  |
| 31-35 | 220 (13.7) | 313 (13.1) | 533 (13.4) |  |
| 36-40 | 194 (12.1) | 267 (11.2) | 461 (11.6) | NS-P>0.05 |
| 41-45 | 141 (8.8) | 204 (8.6) | 345 (8.7) |  |
| 46-50 | 97 (6.1) | 177 (7.4) | 274 (6.9) |  |
| 51-55 | 101 (6.3) | 120 (5.0) | 221 (5.5) |  |
| 56-60 | 63 (3.9) | 96(4.0) | 159 (4.0) |  |
| 61-65 | 48 (3.0) | 92 (3.9) | 140 (3.5) |  |
| 66-70 | 30 (1.9) | 53 (2.2) | 83 (2.1) |  |
| **Marital status** |  |  |  |  |
| Single | 395 (24.6) | 232(9.7) | 627(15.7) |  |
| Married/in relationship | 1064(66.3) | 1580(66.3) | 2644(66.3) |  |
| Widowed/other | 145(9.1) | 570(23.9) | 715(17.9) | P<0.001 |
| **Education†** |  |  |  |  |
| None | 123(7.7) | 531(22.3) | 654(16.4) |  |
| Primary | 643(40.1) | 983(41.3) | 1626(40.8) |  |
| Secondary+ | 826(51.5) | 865(36.3) | 1691(42.4) | P<0.001 |
| **Work status** |  |  |  |  |
| Employed | 1216(75.8) | 1385(58.1) | 2601(65.3) |  |
| Student/unpaid worker | 313(19.5) | 832(34.9) | 1145(28.7) |  |
| Unemployed | 75(4.7) | 165(6.9) | 240(6.0) | P<0.001 |
| **Income per month-** |  |  |  |  |
| **median (IQR)** | 110,000(40,000-300,000) | 60,000 (20,000-200,000) | 100,000 (20,000-200,000) | P<0.01 |
| **Drinks alcohol 3 times/more a week** | 252(15.7) | 81(3.4) | 333(8.4) | P<0.001 |
| **Obese (body mass index >30 kg/m<sup>2</sup>)** | 39(2.4) | 259(10.9) | 298(7.5) | P<0.001 |
| **Has hypertension (blood pressure-systolic ≥140mmHg or diastolic≥90)††** | 441(28.1) | 604(25.9) | 1045(26.8) | NS-P>0.05 |

NB: all the above figures are unweighted

The level of frequent alcohol consumption (3 or more times a week) was 8.4% but it was significantly higher among men (16%) than women (3.4%). Obesity was at 7.5% and it was higher among women (10.9%) than men (2.4%). Hypertension level was at 18.4% and it did not significantly differ by sex.

### Alcohol consumption, hypertension and alcohol consumption across different age groups

Fig 1 shows the trend of frequent alcohol consumption, hypertension and obesity across age groups. The levels of hypertension and frequent alcohol consumption rise with age group and almost at the same average gradient until 46-50 years when the prevalence of hypertension rises higher. The level of obesity reduces with age group to near zero in age group 51-55 when it rises, but at a lower gradient. Values for all the three indicators start at nearly the same level and sharply diverge after 26-30 year age group. A significant test of the gradient of each of the trends in the figure below showed significance (p<0.01).

**Fig 1:**
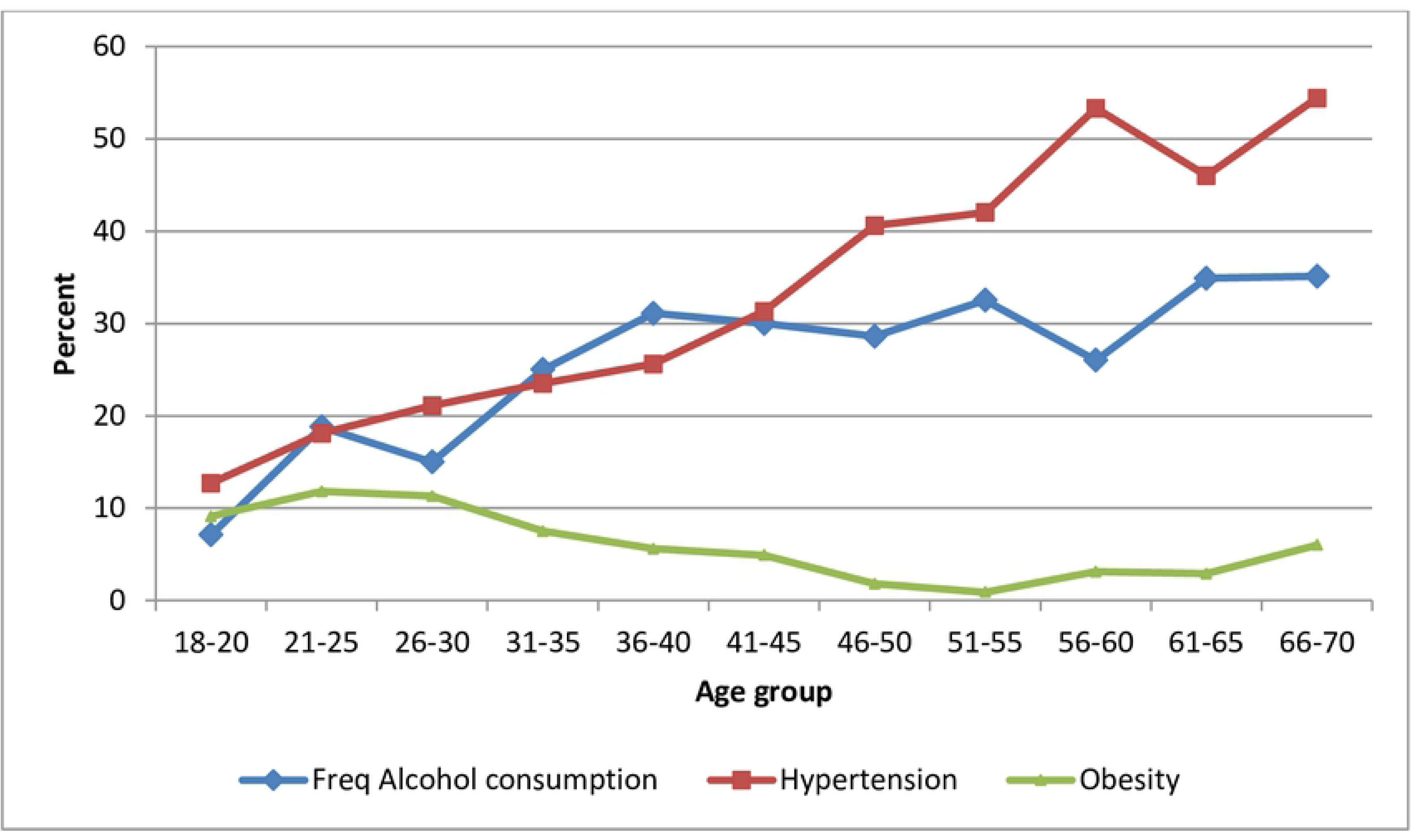
Levels of hypertension, obesity and frequent alcohol consumption across different age groups.

### Effect of alcohol on the Hypertension- age relationship

The prevalence ratios for hypertension rose by age group and this persisted after controlling for frequency of alcohol consumption and other key factors (Table 2). A test of significance of an interaction between frequency of alcohol consumption, hypertension and age group did not yield any significance at 5% level but when age group was replaced with age in single years it was significant. This shows that the relationship between alcohol consumption and hypertension significantly changed by single year rather than by age group.

**Table 2:** Hypertension Prevalence ratios along different age groups and other factors.

| Characteristics | Prevalence<br>N (%) | Un-adjusted<br>Prevalence ratio | Prevalence ratio adjusted<br>for alcohol consumption |
| --- | --- | --- | --- |
| **Age** |  |  |  |
| 18-20 | 53(12.7) | 1.0 | 1.0 |
| 21-25 | 119 (18.1) | 1.43 (1.06-1.93)* | 1.52 (1.12-2.06)** |
| 26-30 | 139(21.1) | 1.67 (1.25-2.23)** | 1.84 (1.36-2.49)*** |
| 31-35 | 123(23.5) | 1.86 (1.38-2.50) *** | 2.10 (1.53-2.86)*** |
| 36-40 | 116(25.6) | 2.02 (1.50-2.72) *** | 2.27(1.66-3.12) *** |
| 41-45 | 104(31.3) | 2.48 (1.84-3.34) *** | 2.80(2.03-3.85) *** |
| 46-50 | 110(40.6) | 3.21 (2.40-4.29) *** | 3.65(2.67-4.99) *** |
| 51-55 | 92(42.0) | 3.32 (2.47-4.46) *** | 3.76(2.73-5.17) *** |
| 56-60 | 82(53.3) | 4.21 (3.14-5.64) *** | 4.71 (3.44-6.46) *** |
| 61-65 | 64(46.0) | 3.64 (2.67-4.96) *** | 4.12 (2.94-5.77) *** |
| 66-70 | 43(54.4) | 4.30 (3.12-5.94) *** | 4.76 (3.38-6.72) *** |
| **Drinks alcohol 3**<br>**times/more a week** |  |  |  |
| 3+ times weekly | 104(32.0) | 1 | 1 |
| <3 times weekly | 292(28.8) | 0.90(0.75-1.08) | 1.05 (0.88-1.26) |
| No/doesn't drink | 649(25.3) | 0.79 (0.67-0.94)** | 1.01 (0.85-1.20) |
| **Sex** |  |  |  |
| Male | 441(28.0) | 1 | 1 |
| Female | 604(25.9) | 0.92 (0.83-1.02) | 0.97 (0.87-1.08) |
| **Marital status** |  |  |  |
| Single | 128(20.9) | 1.0 | 1 |
| Married/in relationship | 675(26.0) | 1.25 (1.05-1.47)* | 0.79 (0.66-0.95)* |
| Widowed/other | 242 (34.8) | 1.67 (1.39-2.01)*** | 0.82 (0.66-1.02) |
| **Work status** |  |  |  |
| Employed | 108(27.9) | 1.0 | -- |
| Self employed | 594(27.5) | 0.98 (0.83-1.17) | -- |
| Student/unpaid worker | 270(24.1) | 0.86 (0.71-1.04) | -- |
| Unemployed | 73(30.9) | 1.11 (0.86-1.42) | -- |
| **Education†** |  |  |  |
| None | 194(30.4) | 1.0 | -- |
| Primary | 429(26.8) | 0.88(0.76-1.02) | -- |
| Secondary+ | 316(24.6) | 0.81(0.70-0.94)** | -- |
| Tertiary | 101 (27.9) | 0.92(0.75-1.13) | -- |
| **Average Income per month** |  |  |  |
| <=500,000 | 252(27.1) | 1 |  |
| >500,000 | 19(28.4) | 1.05 (0.71-1.56) | -- |

†15 refused to provide information. -- left out because Wald test showed $p > 0.1$

Fig 2 shows the ratio of prevalence of hypertension at each age group to that at base age group of 18-20 by frequency of alcohol consumption. It’s evident that after 40 years the hypertension prevalence ratio across age groups among frequent drinkers was persistently lower than that among those who did not take alcohol or drank moderately while trends for those who drank moderately and those who never drank kept a steady rise at almost equal gradient. Beyond 60 years the ratio among the frequent drinkers dropped sharply to 0.8 rose slightly to 2.3.

**Fig 2:**
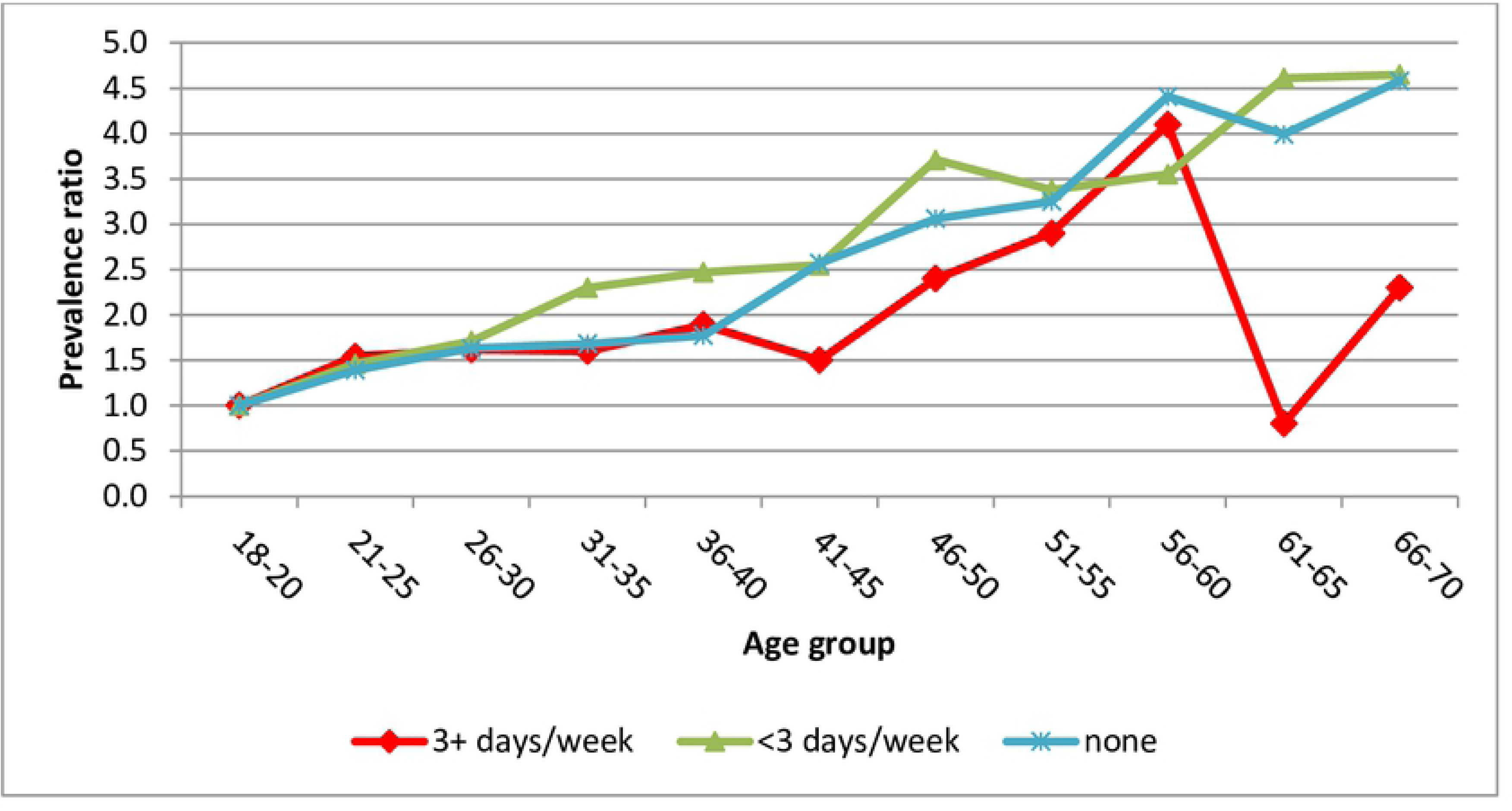
Prevalence ratios for Hypertension across age groups by frequency of alcohol consumption among both men and women.

A closer examination of trends of prevalence ratios across age groups by sex showed that the prevalence of hypertension across age groups is relatively lower among men that don’t take alcohol while it’s the opposite among women (Fig 3). A Wilcoxon’s rank sum test for the difference was significant for men (p=0.03) but not for women (p>0.1). The figure left out those who drank most frequently because they were too few to split by sex across age groups.

**Fig 3:**
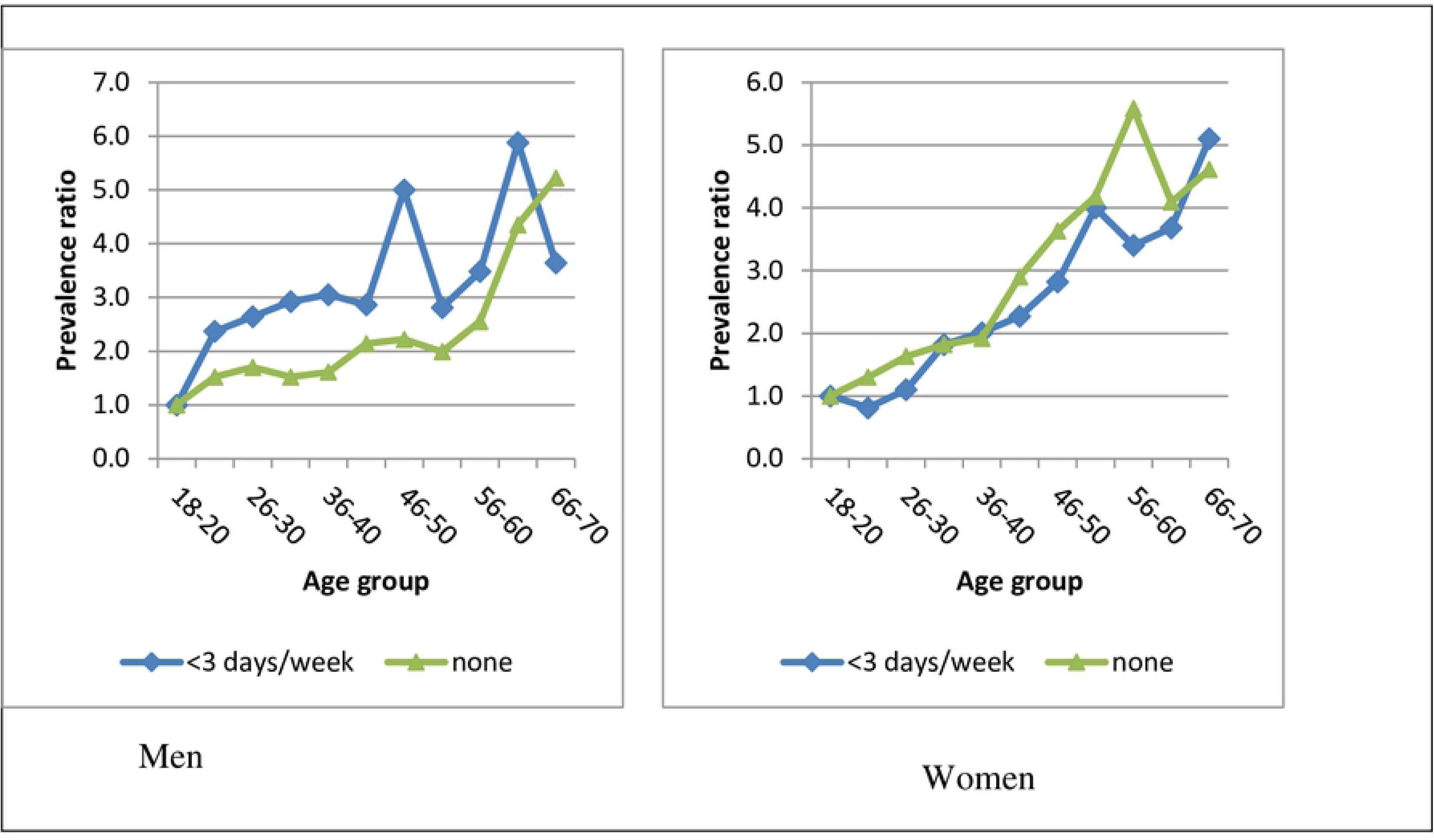
Prevalence ratios for Hypertension across age groups by frequency of alcohol consumption and by sex.

Fig 4 shows fitted lines for prevalence ratios for hypertension at different age groups by alcohol consumption patterns. The figure complements results in Fig 2 and shows an interaction of drinking pattern on hypertension-age group relationship which was not significant on use of age group but significant on use of single year age.

**Fig 4:**
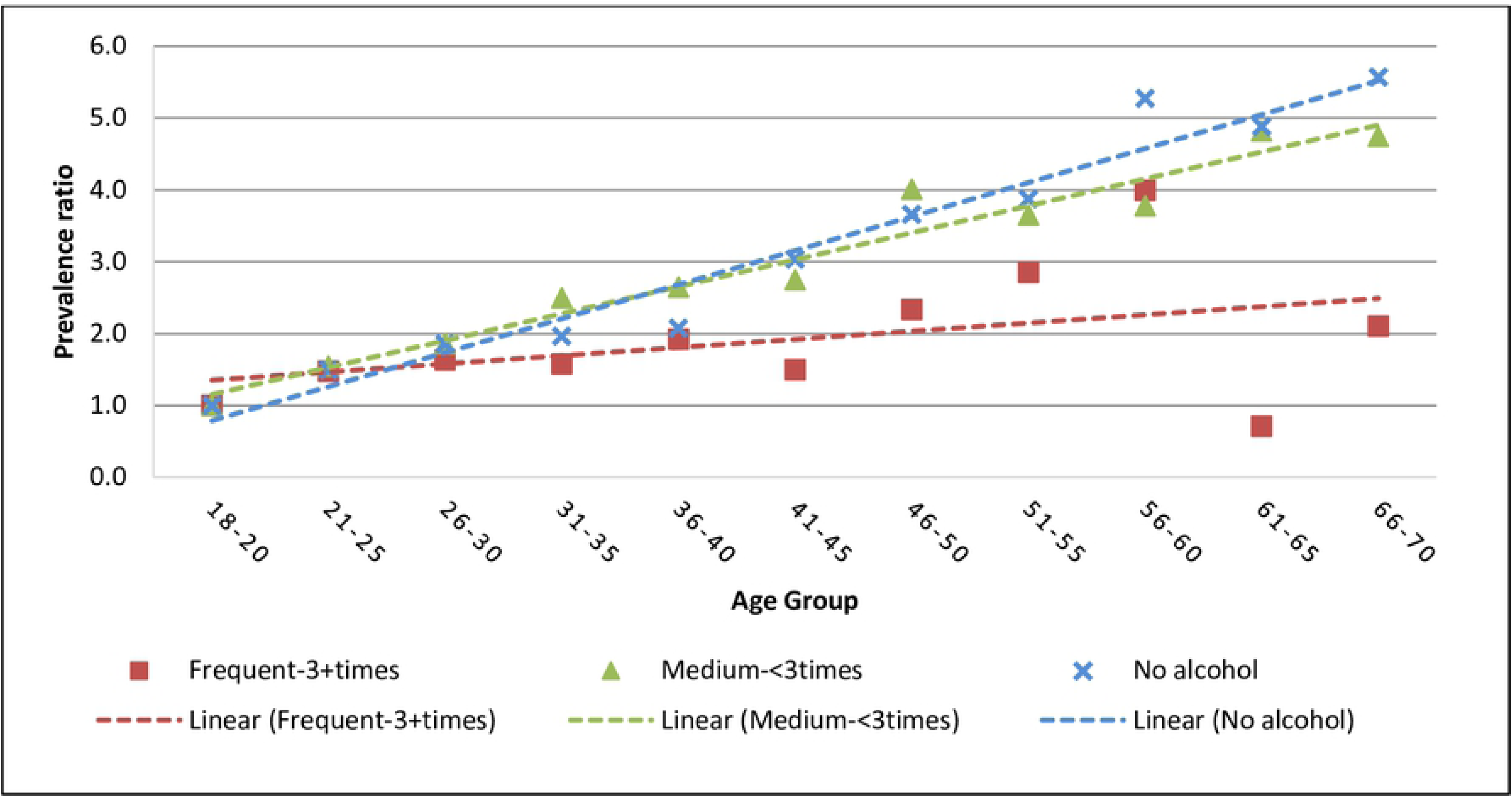
Fitted prevalence ratios for hypertension across different age groups by alcohol consumption pattern.

A Wilcoxon’s signed rank test of significance between the trends for drinking 3+days weekly & <3 days weeks showed a significant difference (p<0.001). The same level of significance was established with comparison of the trends for drinking 3+ weekly & No alcohol. Each of the fitted lines had a statistically significant gradient.

### Obesity

Fig 5 shows the trend of prevalence of obesity across all age groups. The prevalence of obesity among women starts high up from around 18% in the age group 21-25 and declines to around 1% at the age group of 46-50 years while that among men starts low at 3.6% and reduces to 3.1% in the same period. The test for the difference in the two trends using Wilcoxon’s rank sum test shows a p-value of p=0.016.

**Fig 5:**
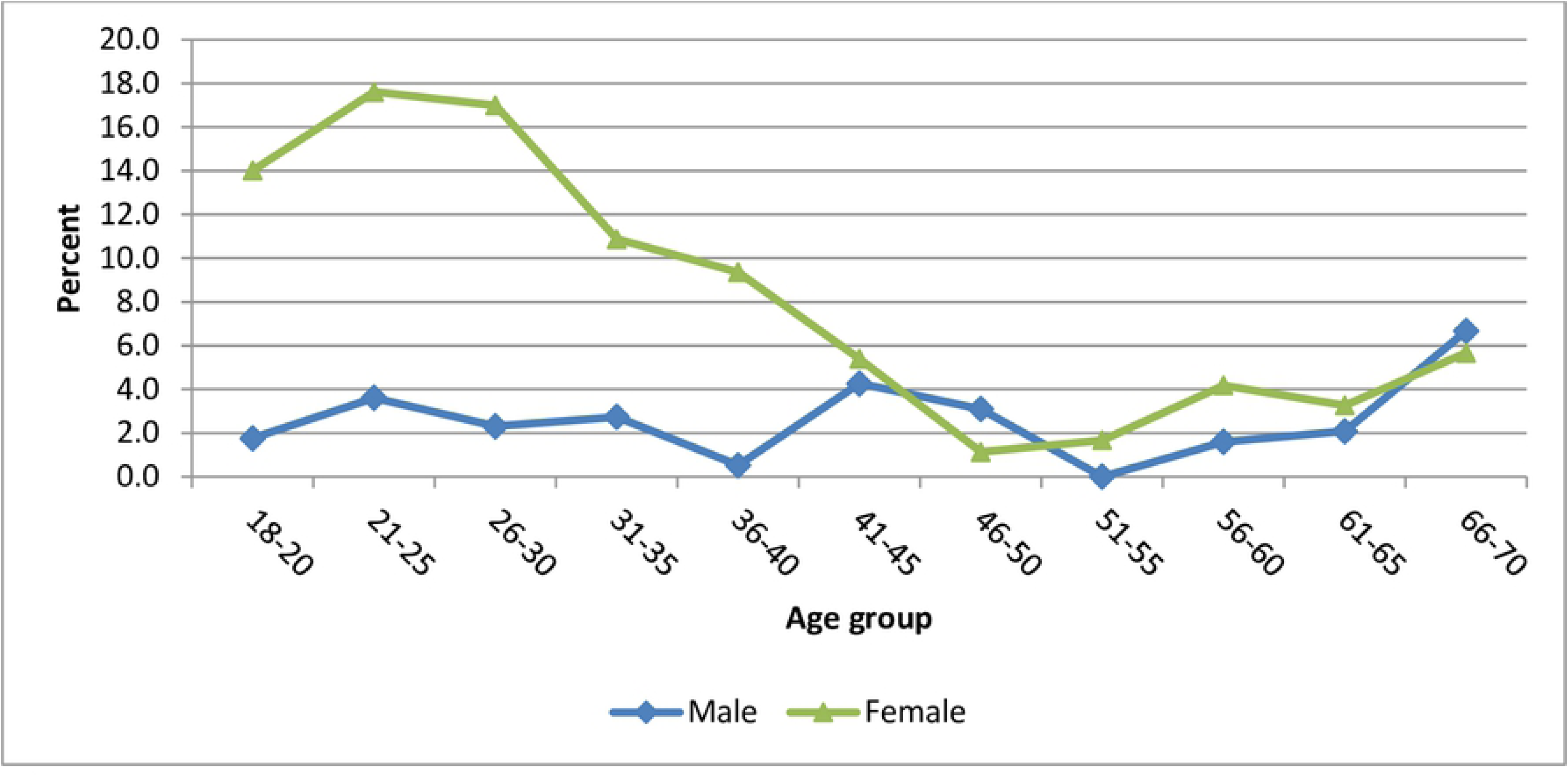
Prevalence of obesity by sex across age groups.

Table 3 shows prevalent ratios for obesity. The prevalence ratios of obesity reduced with increasing age groups even after controlling for drinking patterns and marital status which were significant in the bivariate analysis. A test of interaction with frequency of alcohol consumption did not show any significance hence lack of influence on the obesity-age group relationship.

**Table 3:**
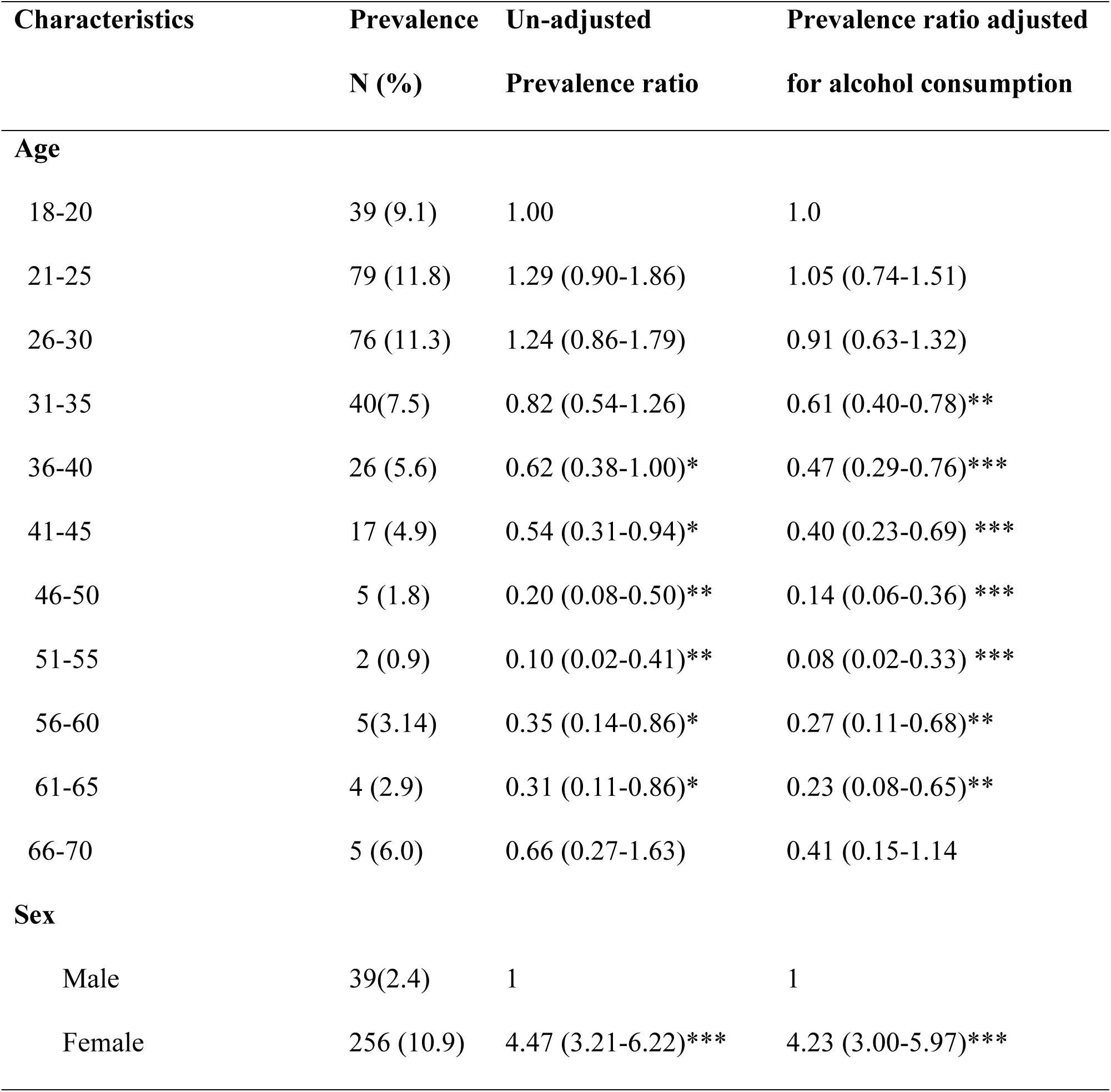
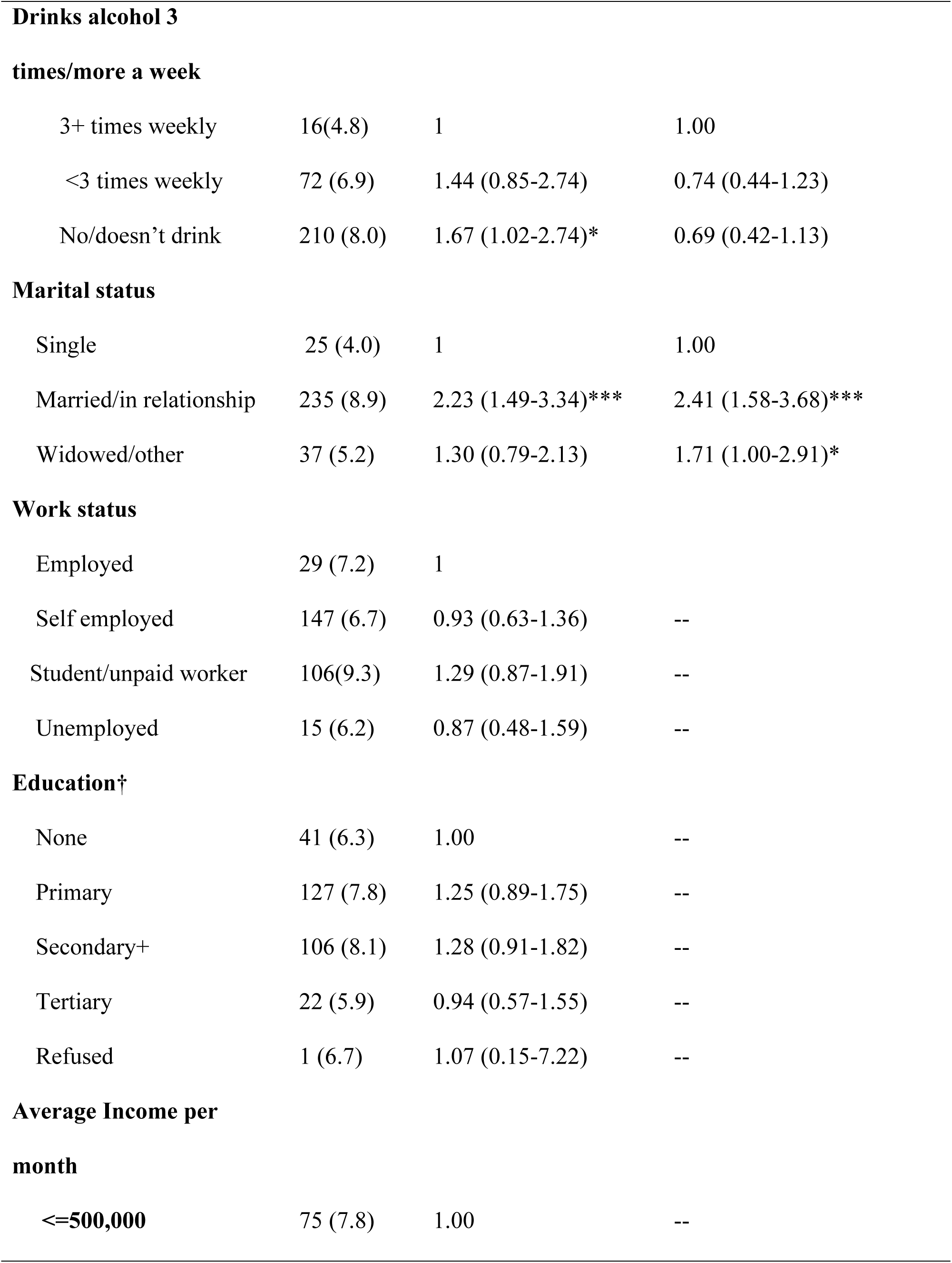
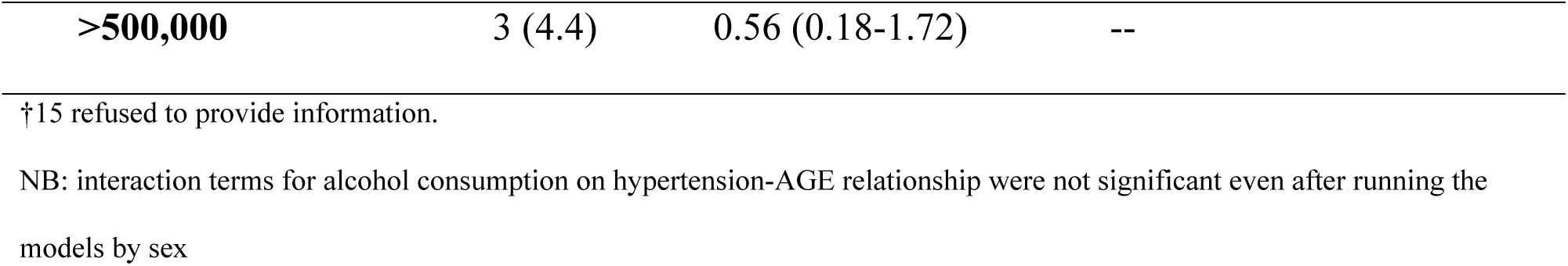
Obesity Prevalence ratios in different age groups and other factors.

Fig 6 shows the trend for obesity levels over age groups by frequency of alcohol consumption. The chart confirms results from table 3. There is a general declining trend of obesity across the age groups and it did not significantly vary by frequency of alcohol consumption.

**Fig 6:**
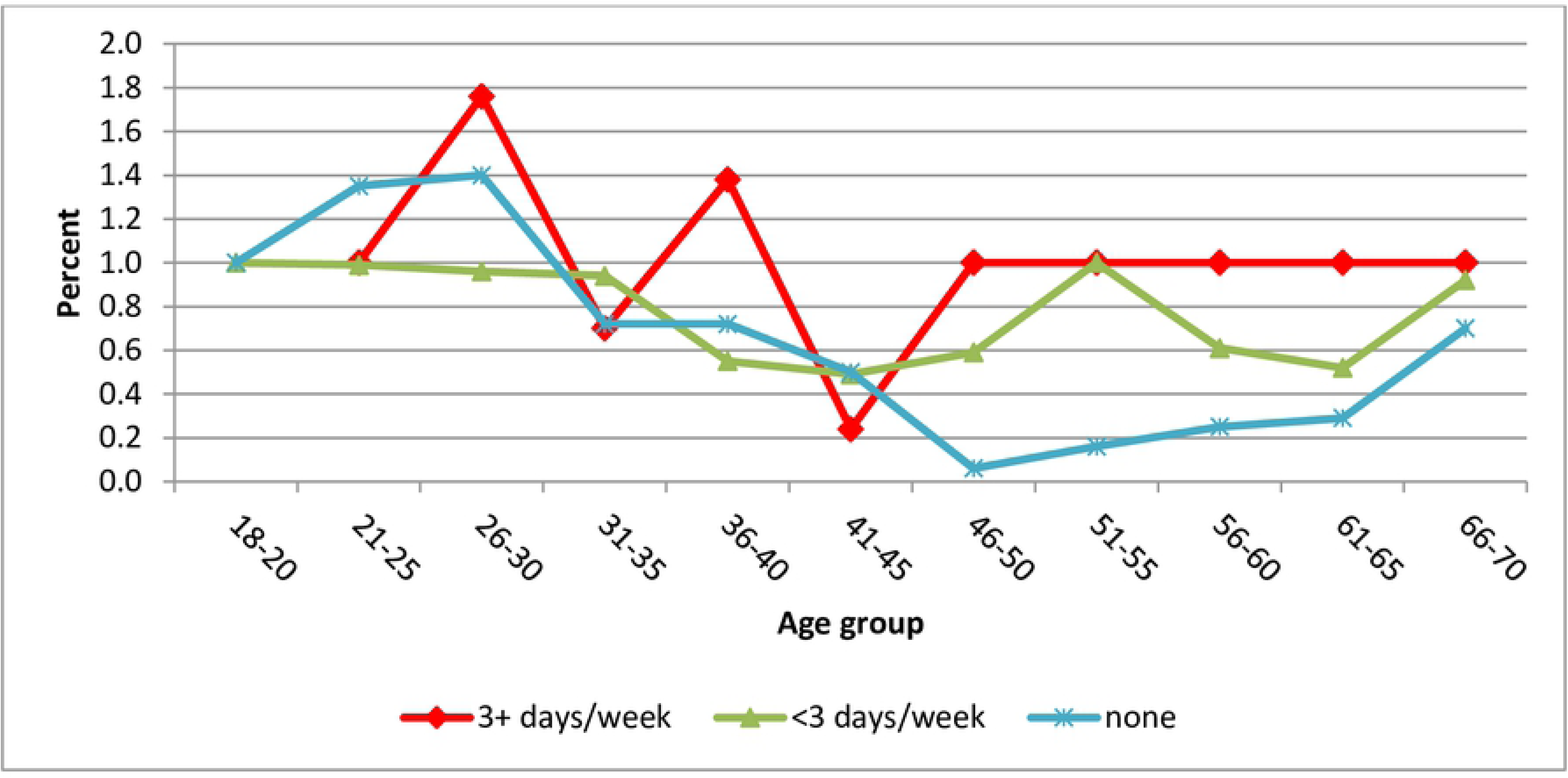
Prevalence ratios for obesity across age groups by frequency of alcohol consumption.

## Discussion

This study explored relationship patterns of alcohol with hypertension and obesity across different age groups. The findings show that the prevalence of frequent alcohol consumption and hypertension follow a nearly similar increasing trend across all age groups below 50 years while the prevalence for obesity follows a downward trend. The relationship between frequent alcohol consumption and hypertension is significant and it changes significantly by age in single years but not by 5 year age groups. The relationship between frequent alcohol consumption and obesity is not significant and does not change across different age groups. Across all age groups the prevalence of hypertension among frequent drinkers is lower than those who either don’t drink or drink less. While among men it’s the non-drinkers that have low prevalence of hypertension it’s the opposite among women.

The increasing trends of frequent alcohol consumption and hypertension across age groups are consistent with several studies in different parts of the world [31] but inconsistent with studies in some other communities[32]. The increasing trend for alcohol consumption can be explained by increased access and ability to buy alcohol which may reduce after 50 years due to change in lifestyle, working environment and social network. Another view could be threshold effect where frequent alcohol consumption exacerbates physiological damages that may also lead to hypertension. However, we noticed that among those who used alcohol frequently, the prevalence ratios rose sharply, dropped and rose again. Future research may examine this pattern more specifically and address other contextual factors not addressed in this study including potential cohort effects, age since alcohol initiation and other potential factors that can impact these findings.

The strong relationship between frequent alcohol consumption and hypertension is reported in many studies [33] and some affirm a causal relationship[34]. However, the change in this relationship with single years and not 5 year age groups is what needs to be investigated further. The prevalence of obesity reduces with age groups but this is more evident among women where it declines sharply between age groups 21-25 and 46-50. This contrasts with studies that show increasing trend of obesity with age group among women in other countries [35, 36] but others show decline with age[37, 38]. While some studies have found a significant positive relationship between alcohol consumption and obesity[39] this study did not find such relationship significant. The negative association between age and obesity can be partially explained by the lower life expectancy observed in Uganda compared to populations examined elsewhere as well as a potential cohort effect of the older participants in this study.

Lower prevalence of hypertension among frequent alcohol consumers compared to the abstainers and infrequent drinkers is an issue that needs further investigation. One probable explanation is the low number of respondents that were frequent alcohol consumers. In subsequent analysis by sex the frequent consumers were left out because of small numbers.

## Conclusion

We conclude, firstly, that the prevalence of frequent alcohol consumption increases across the age groups at almost the same level with prevalence of hypertension until 50 years of age. The frequency of alcohol consumption did not significantly modify the age group-hypertension and obesity-age group relationships but the effect was significant with single years. The prevalence of hypertension among frequent alcohol consumers is lower than that among abstainers and infrequent/mild drinkers. This calls for further research as this is inconsistent with several studies.

While among men it’s the non-drinkers that have lower prevalence of hypertension than the mild drinkers across age groups it’s the opposite among women although the difference is not significant as in the case with the men. More research is needed to identify causes behind this difference.

Programs aimed at reducing hypertension should include messages on abstinence from alcohol consumption especially among men. Priority should also go to older persons.

Frequent alcohol consumption is a key factor in the prevalence of hypertension among most age groups examined, but particularly among those ages 40 and above. Frequent drinking is a modifiable factor that needs to be addressed within clinical practice. Typically, while treatment protocols may include counseling patients on the risk of alcohol, it is not clear whether doctors or health care providers in Uganda have fully embraced or implemented alcohol reduction strategies such as alcohol screening and brief interventions in their treatment of hypertension (is this true). However, this modification to current clinical practice should be explored and investigated further. Moreover, additional research is needed to determine the biological mechanism linking alcohol use to hypertension and what factors exacerbate this association among adults in their 40-50s.

These findings underscore the importance of examining alcohol use in the context of non-communicable diseases in order to determine prevention and intervention strategies.

## Limitations

Our findings should be considered in the context of several limitations. First and foremost, the data are self-reported and as such, inherent biases or lack of knowledge about certain health conditions may have yielded an underestimate of both obesity and hypertension. Moreover, while disclosing alcohol use is not considered a sensitive matter, study participants across settings often under report actual use. Most likely these limitations yield an underestimate of the true association between alcohol use, hypertension and obesity than what has been reported in this study. Moreover, the analyses were based on a cross sectional survey and as such we could not measure the timing and prospective association of the associations between alcohol use and hypertension and obesity which would be of great importance for future prospective cohort studies. Finally, we included several potential confounders in our analyses. However, there may be other important variables that were not considered or available for analyses.

## Ethics

The principal investigator obtained a written permission to use the secondary data from the management of the non-communicable diseases risk factor survey of 2014.

## Acknowledgement

We thank the investigators and sponsors of the Uganda National Study of Non-Communicable Diseases risk factor survey of 2014. We further thank all staff and participants in the study without their good work; we would not prepare this report.

## References

1. Johnson, W., Li, L., Kuh, D., and Hardy, R., How has the age-related process of overweight or obesity development changed over time? Co-ordinated analyses of individual participant data from five United Kingdom birth cohorts. PLoS medicine, 2015. 12(5): p. e1001828.

2. Ren, Q., Su, C., Wang, H., Wang, Z., Du, W., and Zhang, B., Prospective study of optimal obesity index cut-off values for predicting incidence of hypertension in 18–65-year-old chinese adults. PloS one, 2016. 11(3): p. e0148140.

3. Jiang, S.Z., Lu, W., Zong, X.F., Ruan, H.Y., and Liu, Y., Obesity and hypertension. Experimental and therapeutic medicine, 2016. 12(4): p. 2395–2399.

4. Schneider, M., Bradshaw, D., Steyn, K., Norman, R., and Laubscher, R., Poverty and non-communicable diseases in South Africa. Scandinavian journal of public health, 2009. 37(2): p. 176–186.

5. Musinguzi, G. and Nuwaha, F., Prevalence, awareness and control of hypertension in Uganda. PloS one, 2013. 8(4): p. e62236.

6. Guwatudde, D., Mutungi, G., Wesonga, R., Kajjura, R., Kasule, H., Muwonge, J., Ssenono, V., and Bahendeka, S.K., The epidemiology of hypertension in Uganda: findings from the national non-communicable diseases risk factor survey. PloS one, 2015. 10(9): p. e0138991.

7. van de Vijver, S., Akinyi, H., Oti, S., Olajide, A., Agyemang, C., Aboderin, I., and Kyobutungi, C., Status report on hypertension in Africa-Consultative review for the 6^th^ Session of the African Union Conference of Ministers of Health on NCD’s. Pan African Medical Journal, 2014. 16(1).

8. MacMahon, S.W., Blacket, R.B., Macdonald, G.J., and Hall, W., Obesity, alcohol consumption and blood pressure in Australian men and women. The National Heart Foundation of Australia Risk Factor Prevalence Study. Journal of hypertension, 1984. 2(1): p. 85–91.

9. Fuchs, F.D., Chambless, L.E., Whelton, P.K., Nieto, F.J., and Heiss, G., Alcohol consumption and the incidence of hypertension: The Atherosclerosis Risk in Communities Study. Hypertension, 2001. 37(5): p. 1242–1250.

10. Kotwani, P., Kwarisiima, D., Clark, T.D., Kabami, J., Geng, E.H., Jain, V., Chamie, G., Petersen, M.L., Thirumurthy, H., and Kamya, M.R., Epidemiology and awareness of hypertension in a rural Ugandan community: a cross-sectional study. BMC public health, 2013. 13(1): p. 1151.

11. Kayima, J., Nankabirwa, J., Sinabulya, I., Nakibuuka, J., Zhu, X., Rahman, M., Longenecker, C.T., Katamba, A., Mayanja-Kizza, H., and Kamya, M.R., Determinants of hypertension in a young adult Ugandan population in epidemiological transition—the MEPI-CVD survey. BMC public health, 2015. 15(1): p. 830.

12. Lloyd-Jones, D.M., Evans, J.C., and Levy, D., Hypertension in adults across the age spectrum: current outcomes and control in the community. Jama, 2005. 294(4): p. 466–472.

13. Kopelman, P.G., Obesity as a medical problem. Nature, 2000. 404(6778): p. 635.

14. Gillman, M.W., Cook, N.R., Evans, D.A., Rosner, B., and Hennekens, C.H., Relationship of alcohol intake with blood pressure in young adults. Hypertension, 1995. 25(5): p. 1106–1110.

15. Fortmann, S.P., Haskell, W.L., Vranizan, K., Brown, B.W., and Farquhar, J.W., The association of blood pressure and dietary alcohol: differences by age, sex, and estrogen use. American Journal of Epidemiology, 1983. 118(4): p. 497–507.

16. Milon, H., Froment, A., Gaspard, P., Guidollet, J., and Ripoll, J., Alcohol consumption and blood pressure in a French epidemiological study. European heart journal, 1982. 3(suppl_C): p. 59–64.

17. Wakabayashi, I. and Araki, Y., Influences of gender and age on relationships between alcohol drinking and atherosclerotic risk factors. Alcoholism: Clinical and Experimental Research, 2010. 34: p. S54–S60.

18. van Leer, E.M., Seidell, J.C., and Kromhout, D., Differences in the association between alcohol consumption and blood pressure by age, gender, and smoking. Epidemiology, 1994: p. 576–582.

19. Weissfeld, J.L., Johnson, E.H., Brock, B.M., and Hawthorne, V.M., Sex and age interactions in the association between alcohol and blood pressure. American journal of epidemiology, 1988. 128(3): p. 559–569.

20. Adeloye, D.O., Estimating the burden of selected non-communicable diseases in Africa: a systematic review of the evidence. 2015.

21. UBOS and ICF, Uganda Demographic and Health Survey 2016. 2017, UBOS and ICF: Kampala Uganda and Rockville, Maryland USA.

22. MOH, WHO, UNDP, and World Diabetes Foundation, Non-Communicable Disease Risk factor Baseline survey. 2014, Ministry of Health, Government of Uganda: Kampala.

23. WHO, Global status report on alcohol and health 2018. 2019, World Health Organization: Geneva.

24. Klatsky, A.L. and Gunderson, E., Alcohol and hypertension: a review. Journal of the American Society of Hypertension, 2008. 2(5): p. 307–317.

25. Stranges, S., Wu, T., Dorn, J.M., Freudenheim, J.L., Muti, P., Farinaro, E., Russell, M., Nochajski, T.H., and Trevisan, M., Relationship of alcohol drinking pattern to risk of hypertension: a population-based study. Hypertension, 2004. 44(6): p. 813–819.

26. Organization, W.H., Clinical guidelines for the management of hypertension. 2005.

27. WHO. Obesity. 2019 [cited 2019 28th April 2019]; Available from: https://www.who.int/topics/obesity/en/.

28. Martinez, B.A.F., Leotti, V.B., Nunes, L.N., Machado, G., and Corbellini, L.G., Odds Ratio or Prevalence Ratio? An Overview of Reported Statistical Methods and Appropriateness of Interpretations in Cross-sectional Studies with Dichotomous Outcomes in Veterinary Medicine. Frontiers in veterinary science, 2017. 4: p. 193.

29. Blizzard, L. and Hosmer, W., Parameter estimation and goodness-of-fit in log binomial regression. Biometrical Journal, 2006. 48(1): p. 5–22.

30. Cummings, P., Methods for estimating adjusted risk ratios. The Stata Journal, 2009. 9(2): p. 175–196.

31. Kutnikar, J.V., Basavegowda, M., Kokkada, V., and Ashok, N.C., Prevalence of Hypertension and Assessment of “Rule of Halves” in Rural Population of Basavanapura Village, Nanjangud Taluk, South India. Heart India, 2014. 2(4): p. 99.

32. Eigenbrodt, M.L., Mosley Jr, T.H., Hutchinson, R.G., Watson, R.L., Chambless, L.E., and Szklo, M., Alcohol consumption with age: a cross-sectional and longitudinal study of the Atherosclerosis Risk in Communities (ARIC) study, 1987–1995. American journal of epidemiology, 2001. 153(11): p. 1102–1111.

33. Saremi, A., Hanson, R.L., Tulloch-Reid, M., Williams, D.E., and Knowler, W.C., Alcohol consumption predicts hypertension but not diabetes. Journal of studies on alcohol, 2004. 65(2): p. 184–190.

34. Klatsky, A.L., Alcohol and hypertension: does it matter? Yes. Journal of Cardiovascular Risk, 2003. 10(1): p. 21–24.

35. Al-Kandari, Y., Prevalence of obesity in Kuwait and its relation to sociocultural variables. Obesity reviews, 2006. 7(2): p. 147–154.

36. Aliyu, M.H., Luke, S., Kristensen, S., Alio, A.P., and Salihu, H.M., Joint effect of obesity and teenage pregnancy on the risk of preeclampsia: a population-based study. Journal of Adolescent Health, 2010. 46(1): p. 77–82.

37. Clark, D.O. and Gibson, R.C., Race, age, chronic disease, and disability. Minorities, aging, and health, 1997. 1997: p. 107–260.

38. Seidell, J.C. and Visscher, T.L., Body weight and weight change and their health implications for the elderly. European journal of clinical nutrition, 2000. 54(S3): p. S33.

39. Arif, A.A. and Rohrer, J.E., Patterns of alcohol drinking and its association with obesity: data from the Third National Health and Nutrition Examination Survey, 1988–1994. BMC public health, 2005. 5(1): p. 126.

